# Contributions of Site Heterogeneity in Multi-site Brain-Phenotype Predictions

**DOI:** 10.64898/2026.11.01.763540

**Authors:** Guanru Chen, Megan Jones, Ishaan Gadiyar, Theodore D. Satterthwaite, Andrew Chen, Simon Vandekar

**Affiliations:** Vanderbilt University, Nashville, TN, USA; Baylor College of Medicine, Houston, TX, USA; University of Pennsylvania, Philadelphia, PA, USA; Penn Lifespan Informatics and Neuroimaging Center (PennLINC), University of Pennsylvania, Philadelphia, PA, USA; Penn/CHOP Lifespan Brain Institute (LiBI), Philadelphia, PA, USA; Medical University of South Carolina, Charleston, SC, USA; Vanderbilt University Medical Center, Nashville, TN, USA

**Keywords:** Machine learning, prediction accuracy, Pearson correlation, site effects, harmonization, multi-site neuroimaging, cross-validation

## Abstract

Machine learning studies of brain–phenotype associations typically summarize accuracy with a single Pearson correlation, yet in harmonized multi-site data this metric is shaped by training-set size, feature and outcome variability, and true effect heterogeneity. We analyzed structural MRI data from the Reproducible Brain Charts consortium. After preprocessing and quality control, 3486 participants from 9 sites were retained. Three training set compositions were used to understand contributions of site heterogeneity after harmonization: within-site (Local), size-downsampled multi-site (Multi-Down), and full multi-site (Multi-Full). Another site-agnostic strategy (Overall) pooled all sites into a single dataset. All strategies included the target site in the training and testing samples. These analyses separate the contribution of the site effect from that of sample size. Between-site heterogeneity was extensive (age *I*² ≈ 95%). The overall correlation of age prediction (*r* = 0.73) exceeded the within-site meta-analytic estimate (Multi-Full *r* = 0.63). The improvement of correlation in the pooled dataset was driven by a mechanical inflation from the larger outcome variance. At matched sample size, within-site training outperformed harmonized multi-site training. Increasing the multi-site training sample size did not yield a significant advantage over within-site training. The apparent advantage of cross-dataset training thus appears to reflect training-set size rather than a benefit of multi-site composition per se. The same pattern generally held for the weaker brain-p-factor signals. Together, these results indicate that between-site heterogeneity in machine learning brain– phenotype prediction persists after harmonization. We recommend reporting site-level prediction accuracy and feature dispersion and standard deviation of the outcome variable alongside any pooled estimate to improve inter-site and inter-study comparison of machine learning prediction accuracy.

**Key points:**

1. Between-site heterogeneity in machine-learning brain–phenotype prediction persists after harmonization: across nine CovBat-harmonized sites (*N* = 3,486), within-site models outperformed multi-site models at matched training size, and enlarging the multi-site training set did not consistently surpass within-site training.
2. A single pooled correlation masks this heterogeneity: pooling sites overstated within-site accuracy for age (*r* = 0.73 vs. 0.63), largely because pooling widens the outcome’s range.
3. Site-level accuracy tracked imaging-feature variance and outcome variability; we recommend reporting site-level accuracy, feature variance, and outcome standard deviation alongside any pooled estimate.

## 1 Introduction

Machine learning (ML) models trained on brain imaging data are widely used in brain-phenotype (B-P) association studies to generate individualized predictions across a wide range of outcomes from chronological age to dimensions of psychopathology (Marek et al., 2022; More et al., 2023; Spisak et al., 2023). These predictions ideally can support goals that include precision and preventive medicine, early detection of neurodegenerative disease, and identification of individuals at elevated risk for psychiatric conditions before clinical manifestation (More et al., 2023). By jointly modeling distributed, high-dimensional imaging features rather than testing each region in isolation, these models can capture nuanced multivariate patterns and non-linear relationships between brain structure or function and complex phenotypes that are invisible to traditional mass-univariate approaches (More et al., 2023; Spisak et al., 2023). Multi-site imaging data are increasingly used in large-scale brain-phenotype studies. However, simply pooling scans across sites without correction can introduce substantial systematic bias since the site of acquisition can be predicted from the imaging features with high accuracy (Wachinger et al., 2021). Although advanced harmonization methods exist for adjusting for site effects (Chen et al., 2022; Fortin et al., 2017, 2018), heterogeneity arising from combining different sites or comparing across studies of ML B-P analyses remains incompletely characterized. The conditions under which cross-site generalization succeeds or fails are not yet well established (Adkinson et al., 2024; Rosenblatt et al., 2024).

To quantify the strength of B-P associations, ML studies commonly report standardized effect sizes, prominently the Pearson correlation between predicted and observed phenotypic values. Since Pearson correlation summarizes prediction accuracy within a shift and scale of the predicted values, it is considered as a single clean number to be easily interpreted across different studies, methods, or samples (Jones et al., 2025; Schulz et al., 2024). However, the nature of this metric makes it vulnerable to multiple study-design choices that can systematically affect its value.

The magnitude of Pearson’s correlation in ML brain-phenotype predictions is heavily shaped by factors that vary across sites and studies through three distinct mechanisms. First, sample size systematically affects its magnitude. One prior study demonstrated that small sample size inherently produces substantial sample noise by comparing empirical neuroimaging data to simple simulations (Varoquaux, 2018). As multiple ML B-P studies have shown, Pearson correlation stabilizes only with hundreds to thousands of samples, while B-P studies usually typically have less than a few hundreds of participants (Marek et al., 2022; More et al., 2023; Spisak et al., 2023). Second, sites differ in how much the phenotype itself varies. Because Pearson’s *r* normalizes the covariance by the marginal standard deviations of the predicted and observed values, it measures prediction error relative to the spread of the phenotype in the evaluation sample. A site with a narrow age range or a restricted symptom distribution will yield a lower *r* than a site with a wide phenotype range, even if the model’s absolute prediction error is identical in the two sites. This is the classical problem of range restriction, and it means that differences in *r* across sites cannot be interpreted as differences in model quality alone. The same dependence on sample variability has been documented for standardized effect sizes in brain-wide association studies: in a meta-analysis of 63 MRI studies, those with greater variability in the covariate reported systematically larger standardized effect sizes, even though the underlying biological association is not itself altered by study design (Kang et al., 2024). The third and most important factor is effect heterogeneity, in which the true B-P mapping itself differs across sites. As a consequence of these three factors, Pearson’s correlation not only reflects how well a model has captured the brain-phenotype association, but it is also shaped by site-specific properties such as size and variability. While general best-practice guidelines exist for predictive modeling of individual differences in neuroimaging (Scheinost et al., 2019), specific guidance and recommendations on how to interpret and compare reported prediction accuracies across heterogeneous multi-site studies remains limited.

The goal of this study is to understand important contributors to heterogeneity in prediction accuracy of ML brain-phenotype associations across studies and sites. Our novel contributions include designing three complementary training and evaluation strategies to isolate the impact of data composition from sample size. This approach enables us to quantify how inter-site heterogeneity within a multi-site study can influence the model performance. Using data from the Reproducible Brain Charts consortium (Shafiei et al., 2025), we predict chronological age and the general psychopathology factor (p-factor) using Ridge regression and Random Forest under three training strategies — within-site, multi-site at matched sample size, and multi-site at full sample size — that together allow us to distinguish the contributions of sample size, outcome/covariate distribution, and effect heterogeneity. Resolving these contributions supports more meaningful interpretation of reported prediction accuracies. Two comparisons are of particular interest: differences among sites when each is trained and evaluated locally, and differences between within-site and pooled multi-site training. In both cases, an observed difference in correlation may reflect a genuine change in how well the model captures the brain– phenotype mapping, or it may instead reflect differences in sample size, phenotypic range, and cross-site effect heterogeneity that shift the correlation while leaving the underlying mapping unchanged.

## 2 Methods

### 2.1 Data

#### 2.1.1 RBC data and preprocessing

Imaging features and harmonized psychopathology phenotypes were obtained from the Reproducible Brain Charts (RBC release v0.1; RRID:SCR_027837) consortium (Shafiei et al., 2025), an open data resource that integrates five large-scale developmental neuroimaging studies with 9 different sites: the Brazilian High Risk Cohort (BHRC), the Developmental Chinese Color Nest Project (CCNP), the Healthy Brain Network (HBN), the Nathan Kline Institute–Rockland Sample (NKI), and the Philadelphia Neurodevelopmental Cohort (PNC).

Structural images were processed by RBC using sMRIPrep (v0.7.1) and FreeSurfer (v6.0.1; RRID:SCR_001847), which performed intensity non-uniformity correction, skull-stripping, and cortical surface reconstruction. Each scan was manually rated by 2–5 expert raters and assigned a quality determination of Pass, Artifact, or Fail; we retained only scans rated Pass. For the two studies with longitudinal designs (BHRC and NKI), only the baseline session was retained, so that each participant contributes a single observation. These criteria yielded 4,506 participants. Following the data preparation pipeline shared with a companion study (Jones et al., 2025), the analytic sample was restricted to complete cases on age, sex, all four harmonized psychopathology factor scores, and both the Schaefer 300-parcel and DKT gray matter volume estimates, giving *N* = 3,486 across nine sites. This is broader than the present analyses require and was retained for consistency of the harmonized features.

##### Ethics statement

This study is a secondary analysis of de-identified, openly shared dataset from the Reproducible Brain Charts consortium, which are released without a data use agreement. Each contributing study obtained its own ethical approval and written informed consent from participants or their parents or legal guardians; these procedures are detailed in the RBC data resource paper (Shafiei et al., 2025). No new data were collected for the present study.

#### 2.1.2 Outcomes and imaging features

To evaluate site heterogeneity in ML prediction accuracy, we used age and a general psychopathology factor (p-factor) derived in RBC by fitting confirmatory bifactor models jointly across cohorts to produce a common-scale individual-level score (Shafiei et al., 2025). Imaging features were FreeSurfer regional gray matter volume (GMV) averaged within the 300 cortical regions of the Schaefer 17-network parcellation (Schaefer et al., 2018; Yeo et al., 2011). The medial wall label was excluded as non-cortical, and the temporal and frontal pole labels were excluded because they were not returned for all participants, yielding 300 structural features per participant. RBC also provides gray matter volume for the Desikan-Killiany-Tourville parcellation and a range of functional derivatives, but the present analyses use the Schaefer structural features only.

#### 2.1.3 Site harmonization of imaging features

Imaging features in the RBC were harmonized using CovBat with the default settings (Chen et al., 2022) to remove site-related differences in feature mean, variance, and covariance structure. The acquisition site was treated as the batch variable, and age, sex, and the four harmonized psychopathology factor scores were supplied as covariates so that their associations with the imaging features were preserved. Harmonization was applied to imaging features only: chronological age was left in raw form, and the p-factor outcome is harmonized using a bifactor model (Shafiei et al., 2025). No site-level adjustment was applied to either outcome. Notably, residual between-site differences in harmonized imaging features and the phenotype outcomes are both possible contributors to between-site variability in prediction accuracy.

### 2.2 Machine learning models

We used two complementary ML algorithms with contrasting inductive biases — Ridge and Random Forest regression — to predict each outcome (chronological age, p-factor) from the harmonized GMV features. Ridge regression was fitted using the *glmnet* package (RRID:SCR_015505; Friedman et al., 2010) in R (RRID:SCR_001905). The regularization parameter λ was selected on the training data by 10-fold internal cross-validation using *cv.glmnet* (10-fold, reduced to as few as 3 folds when the training set was small) over a designated grid ranging from 10 to 0, retaining the value that minimized held-out mean squared error. Features were standardized to unit variance prior to fitting. Random Forest regression (Breiman, 2001) was implemented using the *ranger* package (RRID:SCR_022521; Wright & Ziegler, 2017). The number of trees, *mtry*, and minimum node size were tuned on the training data via sequential model-based optimization using *tuneRanger*, minimizing out-of-bag prediction error (Probst et al., 2019).

### 2.3 Cross-validation procedure

To distinguish the contributions of training composition and training-set sample size to ML brain-phenotype prediction accuracy, we implemented three training strategies that share a common evaluation procedure but differ in how the training set is constructed. For each of the nine sites in turn, we treated that site as the target and used a 5-fold cross-validation scheme within the target site to define five evaluation folds. In each fold iteration, 20% of the target site’s participants were held out as the evaluation set, while the remaining 80% formed the target-site training portion. The three strategies, defined below, differ in how the target-site portion is combined with — or jointly downsampled together with — data from the eight non-target sites to form the model’s actual training set. In addition to the following three strategies, the data was fitted on the entire sample using the 5-fold cross-validation procedure to obtain a single multi-site estimate of prediction accuracy.

#### 2.3.1 Local

The training set consists of 80% of the target site only — no data from other sites are used. This corresponds to the within-site prediction 5-fold cross-validation setting commonly reported in a single-site B-P association study.

#### 2.3.2 Multi-Full

The training set consists of 80% of the target site combined with all participants from the eight non-target sites. This represents the largest multi-site training set available and reflects the multi-site training setting common in large consortium analyses.

#### 2.3.3 Multi-Down

The training set for the Multi-Down approach is a site-stratified random downsample drawn from the set of all training-eligible participants (the 80% of the target site combined with all participants from the eight non-target sites). The total downsample size was matched to that of the Local training set. Sampling was stratified by site, with each contributing site supplying a number of participants in proportion to its share of the full multi-site sample. This design preserves the relative site composition of Multi-Full while sharing the smaller training-set size of Local.

#### 2.3.4 Overall

In addition to the three site-targeted strategies, we also estimated an overall, site-agnostic prediction by pooling all participants across nine different sites into a single dataset (*N* = 3,486) with standard 5-fold cross-validation.

#### 2.3.5 Repeated splits

The 5-fold partitioning was repeated 100 times with independent random fold assignments to characterize the stability of prediction accuracy across splits, yielding, for each site-strategy combination and for the overall pooled setting a distribution of split-level Pearson correlations between predicted and observed phenotype values. Models were refitted on each split. Across the 100 repeated cross-validation partitions, we computed a split-level Overall Pearson correlation. Within each split, a Pearson correlation was computed separately on each of the five held-out folds and averaged across folds, matching the aggregation used for the three site-targeted strategies. The Overall point estimate reported in Results is the median of these 100 split-level correlations (Chernozhukov et al., 2018), and its 95% interval is the 2.5th–97.5th empirical percentile range of the same 100 values. For site-targeted strategies, the confidence interval was calculated using meta-analytic methods, which will be introduced in the following section.

### 2.4 Rationale and interpretation of the three-strategy design

The comparison between Local and Multi-Down involves equal training set sizes but different compositions; therefore, any differences in evaluation accuracy are due to site-related factors, such as effect heterogeneity and variations in feature and outcome distributions, rather than the size of the training sample. In contrast, the comparison between Multi-Down and Multi-Full holds composition similar while varying training sample size, meaning any differences can be largely attributed to training sample size alone. Finally, comparing Local to Multi-Full confounds both composition and size, and therefore requires extra care in interpretation.

### 2.5 Combining site-level estimates via random-effects meta-analysis

For each site-strategy combination, the median split-level Pearson correlation across the 100 splits served as the site-level summary of prediction accuracy. To combine these nine site-level estimates into a single cross-site meta-analytic estimate per strategy, we conducted a random-effects meta-analysis using the *metacor* function from the *meta* R package (RRID:SCR_019055; Balduzzi et al., 2019). Site-level correlations were Fisher *z*-transformed *z* = atanh(*r*), with the sampling variance of each *z* approximated as 1/(*n* − 3), where *n* is the target site’s sample size. Between-site heterogeneity (*τ*²) was estimated by restricted maximum likelihood (REML), and the meta-analytic *z*-estimate was computed as the inverse-variance-weighted average under the random-effects model. The meta-analytic *z* and its 95% confidence interval were back-transformed to the Pearson correlation scale via *r* = tanh(*z*). We report the random-effects meta-analytic estimate as the primary cross-site summary, and use *I*² (Higgins & Thompson, 2002) to quantify the proportion of total variance attributable to between-site differences in true effect.

Beyond the strategy-specific meta-analyses described above, we additionally included a single multivariate mixed-effects meta-regression, treating the 27 site × strategy Fisher *z* estimates jointly, with strategy as a fixed effect and a site-level random intercept to account for within-site correlation across strategies. Three pre-specified contrasts on this model — Multi-Down vs Local, Multi-Full vs Local, and Multi-Full vs Multi-Down — provided formal pairwise tests of strategy differences. Meta-analyses do not account for correlation among folds; we also performed sensitivity analyses using the Split-HAlf RePeated (SHARP) cross-validation (Zeng et al., 2026), which accounts for correlation among folds (**Supplementary Methods**).

### 2.6 Site-level determinants of prediction accuracy

To examine which site-level properties are associated with the within-site prediction accuracy, we related each site’s Local median Pearson correlation to four site-level summary measures: the standard deviation of the true phenotype, the total variance of imaging features (defined as the trace of the covariance matrix of the imaging features, tr(Cov(*X*))), the standard deviation of the predicted phenotype, and sample size, treating each of the nine sites as a single observation.

## 3 Results

### 3.1 Decomposing site effects on prediction accuracy

To evaluate the contributions of sample size and training composition to the prediction accuracy of ML brain-phenotype associations, we compared cross-validation using the pooled full dataset (Overall) to the three training composition methods (Local, Multi-Down, Multi-Full). All strategies were evaluated for both age and p-factor prediction using ridge regression and random forest regression with structural data in the RBC.

When ridge regression models for age were trained and evaluated on the full RBC sample across all sites, the prediction accuracy was high: *r* = 0.731 (95% CI = [0.728, 0.735]). This overall benchmark exceeded the cross-site random-effects meta-analytic pooled estimates obtained when accuracy was assessed within each site separately: Multi-Full *r* = 0.63 (95% CI = [0.49, 0.74]), Multi-Down *r* = 0.50 (95% CI = [0.35, 0.63]), and Local *r* = 0.58 (95% CI = [0.44, 0.70]; **Figure 1, Supplementary Table 1**). Between-site heterogeneity was extensive within every strategy: Multi-Full *I*² = 95.3% (95% CI = [92.8%, 96.9%]), Multi-Down *I*² = 94.8% (95% CI = [92.1%, 96.6%]), Local *I*² = 95.3% (95% CI= [93.0%, 96.9%]), indicating substantial site-heterogeneity in Pearson estimates (**Supplementary Table 1**).

**Figure 1:**
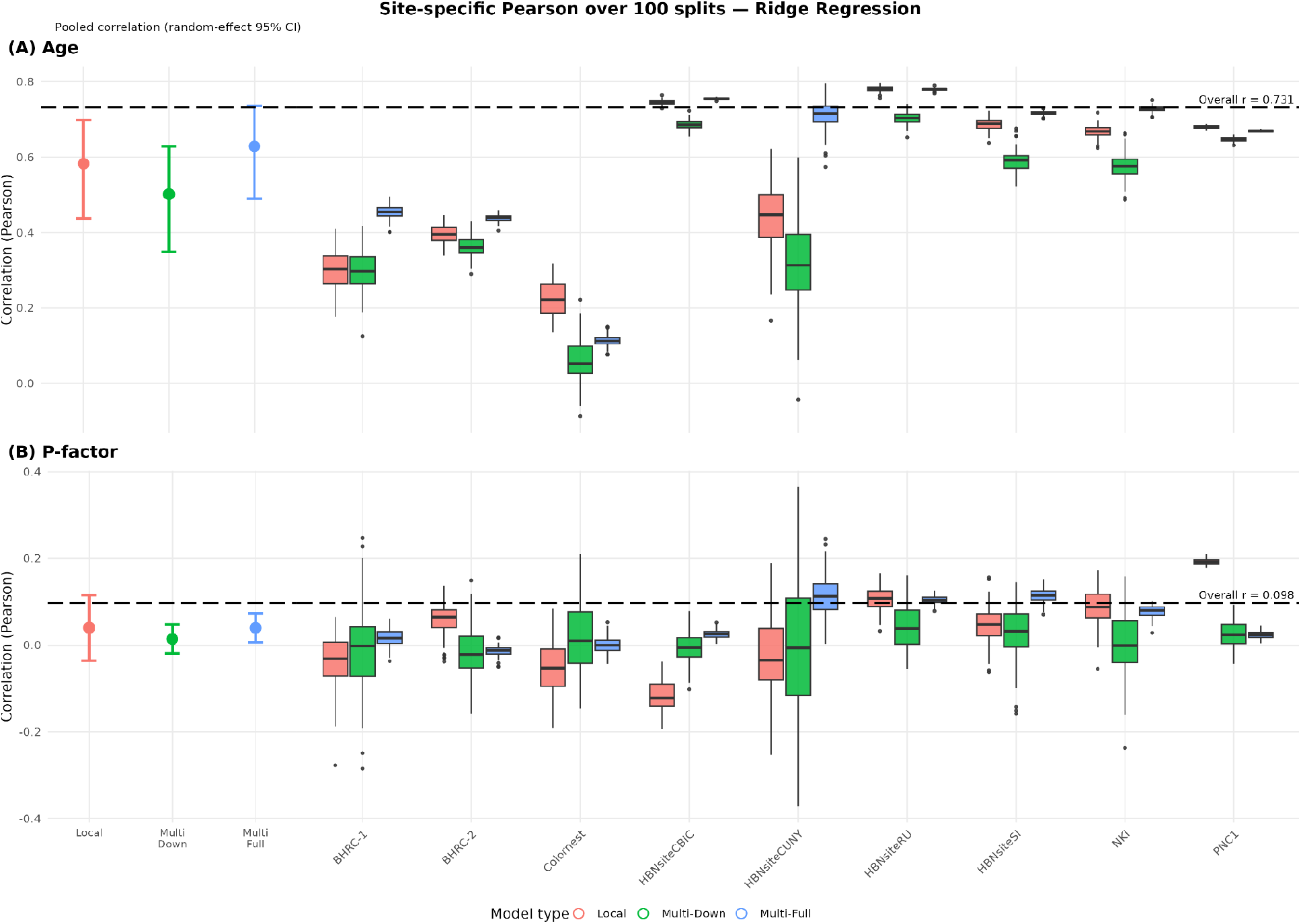
Three modeling approaches were used to evaluate out-of-sample prediction accuracy to quantify the effect of between and within site prediction for different sample sizes for (A) Age and (B) P-factor with site-adjusted imaging data using CovBat. Local Pearson: model trained and tested within site; Multi-Full Pearson: model trained without evaluation fold from target site. Multi-Down Pearson: model trained without evaluation fold from target site and data down-sampled across sites to the size of the target site; dashed lines show the site-agnostic Overall estimate, in which all nine sites are pooled into a single dataset; all methods use 5-fold cross-validation. “Pooled correlation” results show meta-analytic weighted-average estimates across sites. Comparing Local to Multi-Down shows improvements due to within-site prediction; across outcome variables and model types training within-site leads to higher average prediction accuracy. Comparing Multi-Down to Multi-Full shows improvements in across-site prediction due to larger sample sizes; across outcome variables and model types larger samples had higher prediction accuracy. Comparing Multi-Full to Local could be harder to interpret and is addressed in the Discussion. “Site-specific” boxplots show variability of the prediction accuracy across sites over random 100 sample-splits of cross-validation.

Comparing the three cross-validation procedures isolates the contribution of sample size and training composition. Within-site training significantly outperformed multi-site training at matched sample size. Across the nine sites, Local achieved a meta-analytic correlation of 0.58 compared with 0.50 for Multi-Down (Fisher-*z* contrast from the multilevel meta-regression: Δ*z* = 0.098, SE = 0.024, *t*(24) = 4.08, *p* < 0.001). Training on a multi-site composition at a fixed sample size therefore reduced prediction accuracy, even after harmonization. Multi-Full vs. Multi-Down showed a substantial training-size benefit (Δ*z* = 0.116, SE = 0.024, *t*(24) = 4.84, *p* < 0.0001), consistent with the conventional expectation that larger training samples improve predictive accuracy. Notably, Multi-Full and Local yielded statistically indistinguishable performance (Δ*z* = 0.018, SE = 0.024, *t*(24) = 0.76, *p* = 0.45) (**Supplementary Table 2**).

For p-factor, the prediction accuracy of the ridge regression model was consistent with prior research (Marek et al., 2022). The overall estimate was *r* = 0.098 (95% CI= [0.084, 0.114]), while random-effects meta-analytic estimates within sites were smaller: Local *r* = 0.040 (95% CI = [−0.035, 0.116]), Multi-Down *r* = 0.014 (95% CI = [−0.019, 0.048]), and Multi-Full *r* = 0.040 (95% CI = [0.007, 0.073]). There was a high degree of uncertainty in heterogeneity estimates from the meta-analytic models: Multi-Full *I*² = 0% (95% CI = [0%, 64.8%]), Multi-Down *I*² = 0% (95% CI = [0%, 64.8%]), Local *I*² = 82.8% (95% CI = [68.6%, 90.5%]) (**Supplementary Table 3**).

The same three contrasts were examined for p-factor. The site-composition contrast, Local versus Multi-Down, again favored within-site training (Local − Multi-Down: Δ*z* = 0.071, SE = 0.024, *t*(24) = 2.94, *p* = 0.007). The sample-size contrast, Multi-Full versus Multi-Down, was small and not significant (Multi-Full − Multi-Down: Δ*z* = 0.026, SE = 0.024, *t*(24) = 1.07, *p* = 0.296), indicating that a nearly tenfold increase in pooled training data yielded little improvement. Consequently, Multi-Full did not recover the accuracy of Local, although this residual difference did not reach significance (Local − Multi-Full: Δ*z* = 0.045, SE = 0.024, *t*(24) = 1.87, *p* = 0.073; **Supplementary Table 4**). Together, these contrasts suggest that for a low-signal phenotype, site effect is more important than training size: pooling sites may degrade prediction accuracy and increasing training data does not recover it.

The gap between the overall estimate and the site-specific Multi-Full estimate (Δ*r* ≈ 0.10 for age) illustrates the benefit of pooling participants across sites when evaluating prediction accuracy because both models have almost the same training set. To understand how computing prediction accuracy across sites affects accuracy estimates, we plotted the outcome versus predicted values for the two models in two sites (**Figure 2**). When prediction accuracy was re-computed using only the predictions for participants within a targeted site, the within-site correlation dropped substantially: *r* = 0.47 in BHRC-1 and *r* = 0.11 in Colornest (**Figure 2**, right panels) compared to the overall pooled correlation of *r* = 0.73. The discrepancy in the modeling accuracy between the two approaches is driven by the within-site age ranges (BHRC-1 has less span on the x-axis) as well as a difference in the slope of the prediction (Colornest has lower slope). This demonstrates that a high overall Pearson correlation can mask near-zero predictive performance at individual sites and be driven by greater age range from combining multiple sites rather than reflecting genuine improved predictive ability. For random forest models, the results of both age and p-factor are similar but with generally lower prediction accuracies. Local models still outperformed multi-site models in both outcomes, and p-factor prediction results also showed preference for within-site training (**Supplementary Tables 5–9, Supplementary Figure 2**).

**Figure 2:**
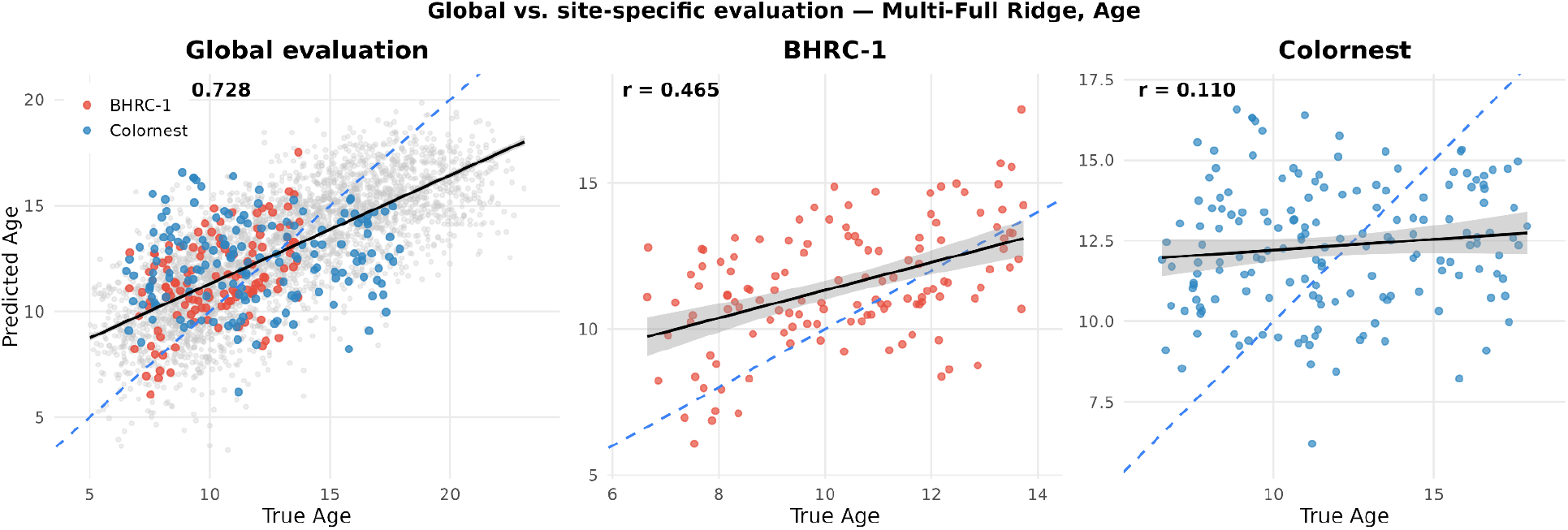
Scatter plots showing the discrepancy between the within-site Pearson correlation and the overall Pearson correlation across sites in two specific sites: BHRC-1 and Colornest, both using the Multi-Full model. The highlighted dots represent individuals from each target site (BHRC-1 in red, Colornest in blue). The y-axis represents the predicted age using Ridge regression and the x-axis represents the true age. Within-site evaluation reveals heterogeneity hidden by global correlation. The same Multi-Full Ridge Regression predictions yield *r* = 0.728 when pooled across all sites (left), but vary widely site-by-site (BHRC-1: *r* = 0.465; Colornest: *r* = 0.110). A high global Pearson correlation can therefore mask near-zero predictive performance at individual sites, which is a direct consequence of site effects. Similar patterns also exist for the Random Forest model, see **Supplementary Figure 1**.

**Figure 3:**
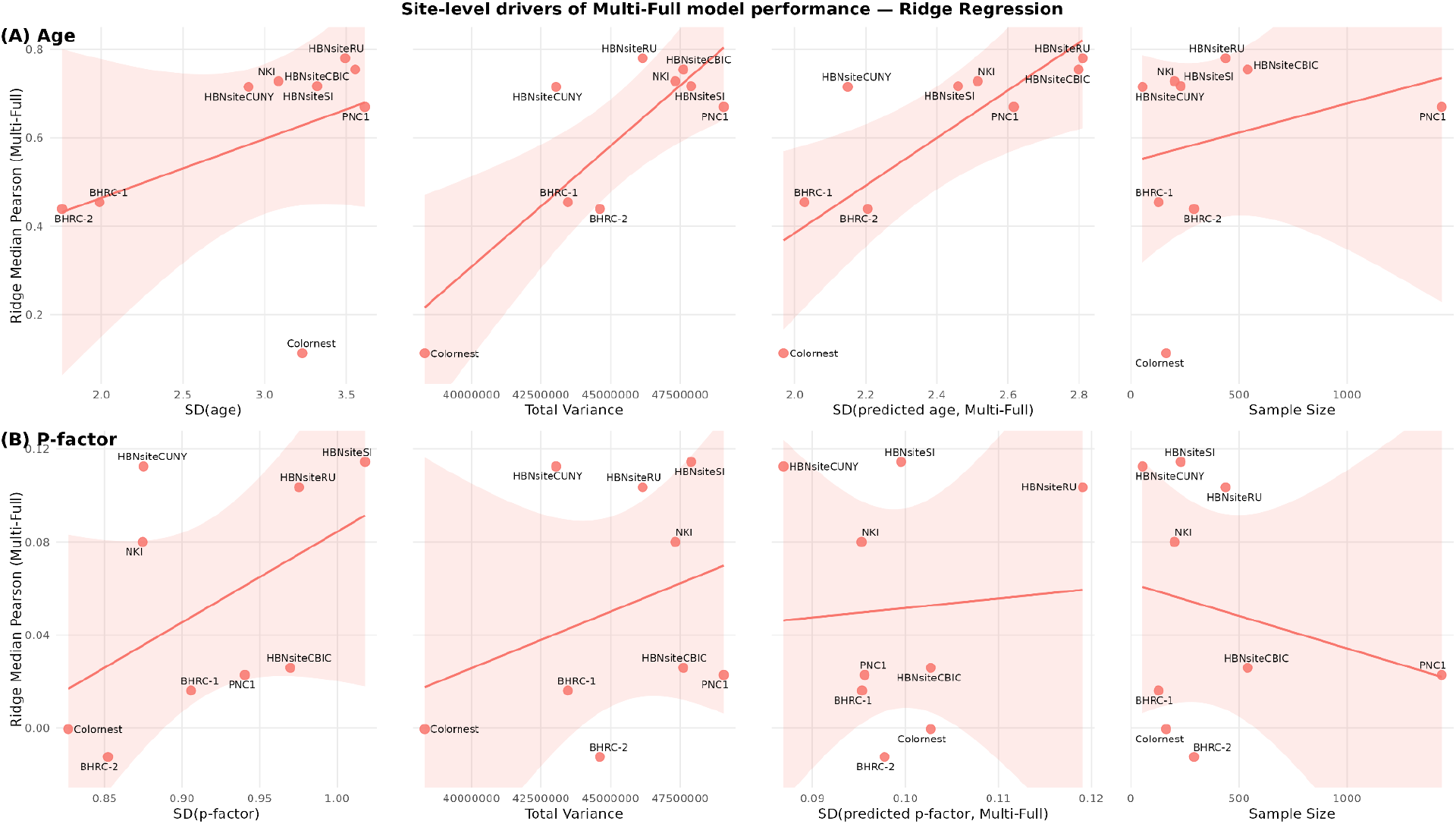
Scatter plots showing marginal association between site-specific Multi-Full Pearson correlation estimate and key site-specific variables for predicting (A) age and (B) P-factor using ridge regression with site-adjusted imaging data using CovBat. Total Variance represents the trace of the covariance matrix of the imaging features for that site.

Our meta-analytic random-effects model did not fully account for dependence among the prediction accuracy estimates arising from shared training samples across sites and shared test samples across strategies. We used SHARP to assess whether the direction and significance were robust after accounting for between-site correlation (**Supplementary Methods**). Effect directions were fully consistent with the original results, with the significance conclusion largely unchanged (Supplementary Table 10). The only difference in significance was the Ridge regression age prediction, where Multi-Full significantly outperformed Local under SHARP. Our sensitivity analyses using SHARP that relax the meta-analysis covariance assumption, show that our main conclusions are robust to the inference method. In particular, our finding that Local has better accuracy than Multi-Down at the matched training sample sizes supports site heterogeneity even after harmonization.

### 3.2 Site-level determinants of prediction accuracy evaluation

To understand how site-level features shape within-site prediction accuracy, we regressed each site’s median Pearson correlation with four summary features for the Multi-Full method: the standard deviation of the outcome, the trace of the imaging-feature covariance matrix tr(Cov(*X*)), which summarizes the total variance of the imaging data, the standard deviation of the predicted values, and the site sample size. Site-level values for all four features, together with the median Pearson correlation under each strategy, are reported in **Supplementary Table 11**. Summary features with Pearson correlation for the Local method are also reported in **Supplementary Figure 3.**

All four summary measures showed positive correlation with the prediction accuracy on age. The total variance of the imaging features, tr(Cov(*X*)) was the strongest determinant (*R*² = 0.68, *p* = 0.006), followed by SD(predicted age) (*R*² = 0.61, *p* = 0.013) and SD(age) (*R*² = 0.17, *p* = 0.27); sample size explained negligible variance (*R*² = 0.06, *p* = 0.51) (**Supplementary Table 12**). The results suggest that sites with larger between-subject heterogeneity in the imaging features and the variability of the phenotype of interest yield higher prediction accuracy. This larger variance is useful insofar as it improves signal-to-noise ratio: under a fixed brain– phenotype mapping, broader total variance tr(Cov(*X*)) widens the range of inputs over which the Pearson correlation is computed. The SD of the predicted values tracks correlation strength closely, but is itself a downstream consequence of the trained mapping rather than an independent driver (see **Supplementary Methods**).

Applied to p-factor, the same analyses produced uniformly weaker associations under the Multi-Full model (*R*² values: SD(p-factor) 0.24, *p* = 0.18; tr(Cov(*X*)) 0.10, *p* = 0.40; sample size 0.05, *p*= 0.55; SD(predicted p-factor) 0.005, *p* = 0.86) (**Supplementary Table 12**). In contrast to age, no single determinant emerged as a dominant driver; the strongest association was with SD(p-factor), but explanatory power was low across all features, consistent with the much weaker brain–phenotype signal at the site level. The corresponding site-level regressions for the Local strategy are reported in **Supplementary Table 13**. For random forest models, the results and figures showed similar messages that both tr(Cov(*X*)) and SD(outcome) are important determinants of prediction accuracies in Multi-Full models (**Supplementary Tables 14 and 15; Supplementary Figures 4 and 5**).

## 4 Discussion

### 4.1 Summary of main findings

We evaluated how three sources of cross-site heterogeneity jointly shape the prediction accuracy in brain-phenotype association studies in harmonized data. First, the conventional pooled Pearson correlation, computed by evaluating one trained model on a held-out test set that mixes participants from all sites, exceeded the random-effects meta-analytic estimate of within-site accuracy under the same trained model. This gap is mechanical: pooling sites broadens the marginal variance of the outcome and inflates Pearson *r* estimates without any improvement in the underlying mapping. Second, at matched training-set sample size, within-site training (Local) significantly outperformed multi-site training (Multi-Down) for both age (Local – Multi-Down: Δ*z* = 0.098, *p* < 0.001) and p-factor (Local − Multi-Down: Δ*z* = 0.071, *p* = 0.007), suggesting that adding other-site data at fixed sample size does not transfer cleanly across the heterogeneity present in the RBC consortium, despite harmonization across sites using CovBat. Third, for age, the gain from a much larger multi-site training set (Local vs. Multi-Full) was almost exactly offset by the cost of cross-site heterogeneity, so that Multi-Full and Local yielded statistically indistinguishable performance. Finally, site-level total variance of imaging features (tr(Cov(*X*))) was the strongest determinant of within-site prediction accuracy for age, although this did not hold in p-factor, which has weaker prediction accuracy. The variance of the imaging features at evaluation enters Pearson correlation mathematically (see **Supplementary Methods**), independent of model training.

### 4.2 Relationship with recent literature

Rosenblatt et al. (2024) showed that prior external validation studies in neuroimaging have routinely been underpowered, combined with significance filtering and publication bias, producing inflated effect-size estimates as a consequence. Our results add a complementary mechanism for accuracy inflation that operates even when sample sizes are adequate: pooling evaluation participants across sites with heterogeneous outcome distributions broadens the variance of the phenotype as well as the variance of the imaging features and mechanically increases Pearson correlation. Under a single fixed trained model, evaluating on the pooled multi-site sample yielded *r* = 0.73, whereas the random-effects meta-analytic summary of the same model’s within-site accuracy was *r* = 0.63, and individual sites ranged as low as *r* = 0.11. Adkinson et al. (2024) demonstrated that brain–phenotype predictions for language abilities and executive function can survive across three diverse, unharmonized developmental datasets, and reported the striking result that for HCPD and HBN, cross-dataset prediction outperformed within-dataset cross-validation. In contrast, our analyses showed the within-site Local strategy significantly outperformed the multi-site Multi-Down strategy at matched training-set size *N*. There are some likely reasons for the discrepancy between our findings: First, Adkinson et al.’s cross-dataset training condition did not control training-set sample size. When one trains on the entire PNC and tests on HBN, the training set is much larger than the within-dataset CV training fold, so apparent advantages of cross-dataset training conflate composition with *N*. Our Multi-Down design holds *N* constant and isolates the composition effect; when matched on *N*, within-site training is the more accurate choice in our data. Second, Adkinson et al. mainly examined cognitive phenotypes like language and executive functions, while we examined age and p-factor.

The brain–outcome mapping for age is strong and stable across development, so although combining sites at the matched sample size degraded within-site accuracy (Local was better than Multi-Down), the loss was small relative to the effect. This loss was offset once the training set was enlarged: Multi-Full and Local were statistically indistinguishable. For the p-factor the same two factors did not balance. The reduction in prediction accuracy when combining sites at the matched sample size was comparable for age and p-factor, but consumed most of the signal (Meta-analytic *r* fell from 0.040 to 0.014 from Local to Multi-Down). Critically, enlarging the training set did not recover it (Multi-Full vs. Multi-Down, Δ*z* = 0.026, *p* = 0.296). Site effects may dominate for a low-signal phenotype in a way it does not for a strong one. One reason is that age is directly observed on a universal scale, whereas the p-factor is a latent score estimated from a measurement model, adding a layer at which the site can enter. Although the RBC harmonized psychopathology to establish measurement comparability, there may not be population equivalence (Hoffmann et al., 2023; Shafiei et al., 2025).

### 4.3 Local-evaluation highlights factors influencing prediction accuracy

Across all three strategies in the age prediction experiment, the discrepancy between the pooled Overall correlation and the within-dataset correlation for the BHRC-1 and Colornest datasets is dramatic. The model itself is not different; marginal variance of the outcome increased the prediction accuracy mechanically by pooling different sites. Notably, these datasets are not outliers on the dimensions one might first suspect: their sample sizes are comparable to other studies (BHRC-1: 128 and Colornest: 163). For BHRC-1, the slope of the line is relatively consistent, but the dataset has a smaller age range that contributes to lower within-site correlation. The one dimension on which Colornest is at the extreme is tr(Cov(*X*)), where it has the smallest value of any site (3.83 × 10^7^ versus 4.30–4.91 × 10^7^ for the others). Although we cannot establish the reason for its outlying value, the practical implication is that the low tr(Cov(*X*)) produces lower range of predictions and lower within-site Pearson correlation under a given trained model. These examples highlight the importance of reporting outcome and feature variance alongside Pearson’s correlation — a low reported correlation from a single site might reflect a model that does not transfer, a population whose features are genuinely less dispersed, or a processing artifact that compresses feature variance.

## 5 Conclusion

Our analyses motivate a set of practical recommendations that build on those of prior papers (Poldrack et al., 2020), organized around the two distinct stages at which multi-site heterogeneity affects reported prediction accuracy: model training and model evaluation.

A larger training sample size is preferable. However, the case for multi-site training is more nuanced. That increasing the size of the training set improves prediction accuracy is a well-established and largely uncontroversial finding; in our data, this is reflected in the substantial advantage of Multi-Full over Multi-Down when training composition is held constant. It is less clear whether one should add other-site data to assemble a larger training sample set. Our Multi-Down vs. Local contrast shows that at a matched training-set size, within-site training outperformed multi-site training for both age and p-factor. For age prediction, the effect of moving from Local to Multi-Full was statistically indistinguishable from zero. Prior work has reported that cross-dataset training can outperform within-dataset cross-validation in certain settings (Adkinson et al., 2024). Such comparisons, however, typically do not hold training-set size constant. The apparent advantage of cross-dataset training may therefore reflect the larger size of the combined training set rather than a benefit of multi-site composition itself. We therefore recommend researchers considering multi-site training explicitly evaluate whether the increase of sample size by pooling different datasets exceeds the cost of cross-site heterogeneity, ideally by reporting both a within-site baseline (Local) and a size-matched multi-site comparison (Multi-Down) alongside the conventional multi-site result.

A high overall correlation does not imply that the model performs comparably on participants from each contributing site. In our data, the overall correlation for age (*r* = 0.73) substantially exceeded the meta-analytic estimate of within-site Multi-Full accuracy (*r* = 0.63), and at the individual site level the same trained model produced correlations ranging from *r* ≈ 0.11 (Colornest) to *r* ≈ 0.79 (HBNsiteRU). Prior methodological guidance has recommended that prediction performance in multi-site data be aggregated at the site level rather than computed over pooled individuals (Dockès et al., 2021). Our results quantified the magnitude of the discrepancy. We therefore recommend reporting site-level prediction accuracy alongside any pooled estimate when using multi-site datasets, so that readers can distinguish a model that performs uniformly across acquisition contexts from one that performs well on average but poorly on specific subgroups. Our site-level analyses identified the total variance of the imaging features, tr(Cov(*X*)), as the strongest exogenous determinant of within-site prediction accuracy under a fixed model (Multi-Full). Under a fixed trained mapping, a site with broader feature variance will yield a larger Pearson *r* at evaluation, all else equal (**Supplementary Methods**). We therefore recommend that authors report the total variance of the imaging features tr(Cov(*X*)) and the standard error of the outcome variable alongside any Pearson-based accuracy estimate, just as one would report sample size.

Our work highlights sample size, outcome variability and imaging predictor variability as key drivers of heterogeneity in ML brain-behavior prediction accuracy. Adherence to our reporting recommendations can help to improve comparisons in prediction accuracy across sites and studies.

**Table 1:** RBC Sample characteristics. *p*-values are from Kruskal-Wallis rank-sum tests and Pearson’s chi-squared test (Sex).

| **Variable** | **N** | **Overall**<br>*N* = 3,486 | **BHRC-1**<br>*N* = 128 | **BHRC-2**<br>*N* = 292 | **Colornest**<br>*N* = 163 | **HBNsiteCBIC**<br>*N* = 540 | **HBNsiteCUNY**<br>*N* = 54 | **HBNsiteRU**<br>*N* = 438 | **HBNsiteSI**<br>*N* = 231 | **NKI**<br>*N* = 202 | **PNC1**<br>*N* = 1,438 | **p-value** |
|---|---|---|---|---|---|---|---|---|---|---|---|---|
| **Age, Mean (SD)** | 3,486 | 12.8 (3.9) | 10.3 (2.0) | 9.9 (1.8) | 11.8 (3.2) | 11.0 (3.6) | 9.9 (2.9) | 11.2 (3.5) | 11.4 (3.3) | 12.9 (3.1) | 15.1 (3.6) | <0.001 |
| **P-factor, Mean (SD)** | 3,486 | −0.14 (1.00) | 0.48 (0.91) | 0.20 (0.85) | −0.44 (0.83) | 0.17 (0.97) | 0.11 (0.88) | 0.19 (0.98) | 0.19 (1.02) | −0.54 (0.87) | −0.46 (0.94) | <0.001 |
| **Sex, n (%)** | 3,486 |  |  |  |  |  |  |  |  |  |  | <0.001 |
| Female |  | 1,600 (46%) | 63 (49%) | 118 (40%) | 90 (55%) | 201 (37%) | 14 (26%) | 158 (36%) | 94 (41%) | 95 (47%) | 767 (53%) |  |
| Male |  | 1,886 (54%) | 65 (51%) | 174 (60%) | 73 (45%) | 339 (63%) | 40 (74%) | 280 (64%) | 137 (59%) | 107 (53%) | 671 (47%) |  |
| **Euler number, Mean (SD)** | 3,486 | −110 (69) | −91 (32) | −73 (23) | −48 (22) | −86 (48) | −87 (44) | −209 (105) | −70 (37) | −83 (37) | −117 (47) | <0.001 |

**Table 2:** Summary of the training strategies. The three site-targeted strategies are evaluated on identical held-out participants and differ only in how the training set is constructed. Overall is not site-targeted and treats the consortium as a single dataset.

| **Strategy** | **Test data and size** | **Training data and size** |
|---|---|---|
| **Local** | Site *k*, held-out fold (20% of site *k*) | 80% of site *k* only |
| **Multi-Full** | Site *k*, held-out fold (20% of site *k*) | 80% of site *k* + all participants from 8 non-target sites |
| **Multi-Down** | Site *k*, held-out fold (20% of site *k*) | 80% of site *k* + 8 non-target sites, site-stratified proportionally downsampled, size matched to Local |
| **Overall** | Held-out fold, all sites mixed (20% of *N* = 3,486) | All 9 sites, no target designation (80% of *N* = 3,486) |

## Data and Code Availability

All imaging features and harmonized phenotypes analyzed in this study are openly available from the Reproducible Brain Charts consortium (RBC release v0.1; Shafiei et al., 2025) at https://reprobrainchart.github.io/ and are shared without a data use agreement. All analysis code is available at https://github.com/guanruchen-vu/Site-effect-RBC.

## Author Contributions

G.C.: Conceptualization, Methodology, Software, Formal analysis, Investigation, Visualization, Writing – original draft, Writing – review & editing. M.J.: Methodology, Software, Writing – review & editing. I.G.: Methodology, Software, Writing – review & editing. T.D.S.: Resources, Funding acquisition, Writing – review & editing. A.C.: Writing – review & editing. S.N.V.: Conceptualization, Methodology, Supervision, Funding acquisition, Writing – review & editing.

## Acknowledgments

This research was supported by National Institutes of Health grants R01MH123563 (to S.N.V.) and R01MH120482 (to T.D.S.).

