## Supplementary Material for "Contributions of Site Heterogeneity in Multi-site Brain-Phenotype Predictions"

### Supplementary Methods

#### Mathematical decomposition of Pearson’s correlation

Pearson’s correlation for machine learning models can be expressed for a training and testing dataset as

$$r = \frac{cov(y_{te}, \hat{\mu}_{tr}(x_{te}))}{sd(y_{te})sd(\hat{\mu}_{tr}(x_{te}))} \ ,$$

where the subscript identifies the dataset used for estimation. The variance of the phenotype and brain imaging data contribute to the prediction accuracy, which can be approximately described using the delta method, assuming the training model, $\hat{\mu}_{tr}$, is fixed

$$cov(y_{te}, \hat{\mu}_{tr}(x_{te})) \approx cov(y_{te}, \frac{\partial}{\partial x}\hat{\mu}_{tr}|_{E(x)} x_{te}) = cov(y_{te}, x_{te})\frac{\partial}{\partial x}\hat{\mu}_{tr}|_{E(x)}$$

$$var(\hat{\mu}_{tr}(x_{te})) \approx \frac{\partial}{\partial x}\hat{\mu}_{tr}|_{E(x)}^{T} var(x_{te})\frac{\partial}{\partial x}\hat{\mu}_{tr}|_{E(x)} \ .$$

These formulas illustrate that the variance of the imaging data contributes to increases in the correlation accuracy insofar as it increases the covariance of the imaging data with the phenotype. In the empirical results (Section 3), we saw that between-site differences in the variance of the imaging data were associated with increased accuracy in prediction, possibly due to differences in imaging quality across sites.

#### SHARP-based sensitivity analysis of strategy comparisons

**Design and contrasts.** We adapted SHARP (Zeng et al., 2026) to assess dependence arising from overlapping training samples across sites and shared test participants across strategies. For each outcome and algorithm, we repeated a joint, site-stratified split into disjoint halves A and B $J$ = 60 times. Each site contributed equally sized halves; one randomly chosen participant was omitted when its eligible count was odd. All training and tuning remained within the assigned half. Five-fold cross-validation used the same target-site test folds for all three strategies. Local used the remaining target-site folds, Multi-Full added the other sites in that half, and Multi-Down proportionally subsampled the Full training pool to match the Local training size.

For site $k$ = 1, …, 9, let $r_{jhkf}^{(s)}$ denote the fold-specific correlation for strategy $s$. With fixed weights based on eligible full-sample counts $n_k$, define

$$z_{jhk}^{(s)} = atanh[\frac{1}{5}\sum_{f=1}^{5} r_{jhkf}^{(s)}], \quad w_k = \frac{n_k - 3}{\sum_{\ell=1}^{9}(n_\ell - 3)},$$

$$D_{jh} = \sum_{k=1}^{9} w_k[z_{jhk}^{(a)} - z_{jhk}^{(b)}].$$

We tested Full − Local, Down − Local, and Full − Down. Each $D_{jh}$ combines paired strategy differences across all sites before inference: its variance is $\boldsymbol{w}^{T}\mathrm{Cov}(\boldsymbol{\delta}_{jh})\boldsymbol{w}$, where $\boldsymbol{\delta}_{jh}$ contains the site-specific differences. Thus, site-level estimates are not treated as independent observations. The target $\mu$ = E($D_{jh}$) concerns this fixed-weight, half-sample design, rather than the original random-effects pooled estimates.

**Working covariance and estimator.** Stack $\boldsymbol{D} = (D_{1A}, …, D_{JA}, D_{1B}, …, D_{JB})^{T}$ and assume $\boldsymbol{D} \sim N(\mu\mathbf{1}, \Sigma)$, with

$$Var(D_{jh}) = \sigma^2, \quad Cov(D_{jA}, D_{jB}) = 0,$$

$$Cov(D_{jh}, D_{j'h'}) = \rho\sigma^2 \quad \text{for } j \neq j'.$$

Disjoint halves motivate zero within-pair covariance under independent participant sampling; a common $\rho$ represents dependence across repeated partitions. Equal variances and this correlation structure are working assumptions. Writing $\Sigma = \sigma^2 R$, the eigenvalues of $R$ are 1, $1 - 2\rho$, and $c = 1 + 2(J - 1)\rho$, with multiplicities $J$, $J$ − 1, and 1. Because $\Sigma\mathbf{1} = \sigma^2 c\mathbf{1}$,

$$\hat{\mu} = \frac{1^T\Sigma^{-1}D}{1^T\Sigma^{-1}1} = \bar{D} = \frac{1}{2J}\sum_{j=1}^{J}(D_{jA} + D_{jB}),$$

$$Var(\bar{D}) = \frac{1}{4J^2}1^T\Sigma 1 = \sigma^2[\frac{1}{2J} + \frac{J-1}{J}\rho].$$

**Null estimation.** For $H_0$: $\mu = \mu_0$, let $u_j = (D_{jA} - D_{jB})/\sqrt{2}$, $v_j = (D_{jA} + D_{jB})/\sqrt{2}$, $A_0 = \sum_j u_j^2$, and $B_0 = \sum_j (v_j - \bar{v})^2$, where $\bar{v}$ is the mean of the $v_j$. The eigendecomposition yields

$$Q(\rho;\ \mu_0) = A_0 + \frac{B_0}{1-2\rho} + \frac{2J(\bar{D}-\mu_0)^2}{c}.$$

The negative Gaussian log-likelihood, up to constants, is

$$-\ell = J\,log\,\sigma^2 + \frac{1}{2}(J - 1)log(1 - 2\rho) + \frac{1}{2}log\,c + \frac{Q}{2\sigma^2}.$$

Differentiation gives $\hat{\sigma}^2(\rho;\ \mu_0) = Q/(2J)$. Substitution into the negative log-likelihood above therefore gives

$$\hat{\rho}_0 = arg\,min_{\rho}\{J\,log[\frac{Q(\rho;\ \mu_0)}{2J}] + \frac{1}{2}(J - 1)log(1 - 2\rho) + \frac{1}{2}log\,c\},$$
$$\hat{\sigma}_0^{\ 2} = \frac{Q(\hat{\rho}_0;\ \mu_0)}{2J}.$$

We optimized over $0 \le \rho \le 1/2 - 10^{-8}$, checking interior and boundary minima.

**Score inference.** With $\hat{\Sigma}_0 = \hat{\sigma}_0^{\ 2}R(\hat{\rho}_0)$, the null score and information are $U_0 = 1^T\hat{\Sigma}_0^{\ -1}(D - \mu_0 1)$ and $I_0 = 1^T\hat{\Sigma}_0^{\ -1}1$. The constant row sum then gives

$$Z_{score}(\mu_0) = \frac{U_0}{\sqrt{I_0}}$$
$$= \frac{\bar{D} - \mu_0}{\sqrt{\hat{\sigma}_0^{\ 2}[\frac{1}{2J} + \frac{J-1}{J}\hat{\rho}_0]}}$$

Using the SHARP standard-normal reference approximation, the two-sided test of $\mu = 0$ and marginal 95% confidence set are

$$p = 2\Phi(-\ |Z_{score}(0)|),$$
$$C_{0.95} = \{\mu_0:\ |Z_{score}(\mu_0)| \le \Phi^{-1}(0.975)\}.$$

Nuisance parameters were re-estimated for each candidate $\mu_0$. Holm correction covered six comparisons per algorithm (three contrasts × two outcomes); confidence sets remained marginal. This multisite adaptation assumes independent participants and the working covariance above. Upstream CovBat was not re-estimated within halves, so harmonization uncertainty was not propagated. Agreement with the primary analysis supports robustness of the conclusions, not validating its covariance assumptions.

### Supplementary Tables

**Supplementary Table 1:** Ridge meta-analysis of within-site Pearson correlations — chronological age. Random- and fixed-effects pooled estimates across the nine sites for each training strategy, with between-site heterogeneity ($\tau^2$, $I^2$).

| Outcome | Strategy | Pooled r (Random) | Pooled r (Fixed) | Random-effects z | Fixed-effects z | $\tau^2$ | $I^2$ [95% CI] |
|---|---|---|---|---|---|---|---|
| Age | Multi-Full | 0.628 [0.490, 0.736] | 0.669 [0.650, 0.687] | 0.739 | 0.808 | 0.0908 | 95.3% [92.8%, 96.9%] |

| Outcome | Strategy | Pooled r (Random) | Pooled r (Fixed) | Random-effects z | Fixed-effects z | τ² | I² [95% CI] |
|---|---|---|---|---|---|---|---|
| Age | Multi-Down | 0.502 [0.349, 0.628] | 0.599 [0.578, 0.620] | 0.551 | 0.692 | 0.0764 | 94.8% [92.1%, 96.6%] |
| Age | Local | 0.582 [0.437, 0.698] | 0.659 [0.639, 0.677] | 0.666 | 0.790 | 0.0860 | 95.3% [93.0%, 96.9%] |

**Supplementary Table 2:** Ridge pairwise strategy contrasts — chronological age. Fisher-z differences between strategies from the multi-level meta-regression (site-level random intercept).

| Outcome | Contrast | Δz | SE (Δz) | t | df | 95% CI (z) | p-value |
|---|---|---|---|---|---|---|---|
| Age | Multi-Down − Local | -0.098 | 0.024 | -4.08 | 24 | [-0.148, -0.049] | 4.29e-04 |
| Age | Multi-Full − Local | 0.018 | 0.024 | 0.76 | 24 | [-0.031, 0.068] | 0.454 |
| Age | Multi-Full − Multi-Down | 0.116 | 0.024 | 4.84 | 24 | [0.067, 0.166] | 6.21e-05 |

**Supplementary Table 3:** Ridge meta-analysis of within-site Pearson correlations — p-factor. Columns as in Supplementary Table 1.

| Outcome | Strategy | Pooled r (Random) | Pooled r (Fixed) | Random-effects z | Fixed-effects z | τ² | I² [95% CI] |
|---|---|---|---|---|---|---|---|
| P-factor | Multi-Full | 0.040 [0.007, 0.073] | 0.040 [0.007, 0.073] | 0.040 | 0.040 | 0.0000 | 0.0% [0.0%, 64.8%] |
| P-factor | Multi-Down | 0.014 [-0.019, 0.048] | 0.014 [-0.019, 0.048] | 0.014 | 0.014 | 0.0000 | 0.0% [0.0%, 64.8%] |
| P-factor | Local | 0.040 [-0.035, 0.116] | 0.085 [0.052, 0.118] | 0.040 | 0.085 | 0.0090 | 82.8% [68.6%, 90.5%] |

**Supplementary Table 4:** Ridge pairwise strategy contrasts — p-factor. Columns as in Supplementary Table 2.

| Outcome | Contrast | Δz | SE (Δz) | t | df | 95% CI (z) | p-value |
|---|---|---|---|---|---|---|---|
| P-factor | Multi-Down − Local | -0.071 | 0.024 | -2.94 | 24 | [-0.118, -0.024] | 0.007 |
| P-factor | Multi-Full − Local | -0.045 | 0.024 | -1.87 | 24 | [-0.092, 0.002] | 0.073 |
| P-factor | Multi-Full − Multi-Down | 0.026 | 0.024 | 1.07 | 24 | [-0.021, 0.073] | 0.296 |

**Supplementary Table 5:** Random Forest meta-analysis of within-site Pearson correlations — chronological age. Columns as in Supplementary Table 1.

| Outcome | Strategy | Pooled r (Random) | Pooled r (Fixed) | Random-effects z | Fixed-effects z | τ² | I² [95% CI] |
|---|---|---|---|---|---|---|---|
| Age | Multi-Full | 0.464 [0.327, 0.581] | 0.529 [0.504, 0.552] | 0.502 | 0.588 | 0.0565 | 92.7% [88.4%, 95.4%] |
| Age | Multi-Down | 0.371 [0.246, 0.484] | 0.466 [0.439, 0.491] | 0.390 | 0.505 | 0.0396 | 90.8% [84.8%, 94.4%] |
| Age | Local | 0.463 [0.320, 0.586] | 0.558 [0.534, 0.580] | 0.502 | 0.630 | 0.0622 | 93.1% [89.1%, 95.7%] |

**Supplementary Table 6:** Random Forest meta-analysis of within-site Pearson correlations — p-factor. Columns as in Supplementary Table 1.

| Outcome | Strategy | Pooled r (Random) | Pooled r (Fixed) | Random-effects z | Fixed-effects z | τ² | I² [95% CI] |
|---|---|---|---|---|---|---|---|
| P-factor | Multi-Full | -0.025 [-0.067, 0.018] | -0.015 [-0.048, 0.018] | -0.025 | -0.015 | 0.0010 | 21.8% [0.0%, 62.7%] |
| P-factor | Multi-Down | 0.002 [-0.031, 0.036] | 0.002 [-0.031, 0.036] | 0.002 | 0.002 | 0.0000 | 0.0% [0.0%, 64.8%] |
| P-factor | Local | 0.052 [-0.025, 0.129] | 0.103 [0.070, 0.136] | 0.052 | 0.103 | 0.0096 | 84.7% [72.8%, 91.4%] |

**Supplementary Table 7:** Random Forest pairwise strategy contrasts — chronological age. Columns as in Supplementary Table 2.

| Outcome | Contrast | Δz | SE (Δz) | t | df | 95% CI (z) | p-value |
|---|---|---|---|---|---|---|---|
| Age | Multi-Down − Local | -0.125 | 0.024 | -5.19 | 24 | [-0.174, -0.075] | 2.55e-05 |
| Age | Multi-Full − Local | -0.041 | 0.024 | -1.72 | 24 | [-0.091, 0.008] | 0.098 |
| Age | Multi-Full − Multi-Down | 0.083 | 0.024 | 3.47 | 24 | [0.034, 0.133] | 0.002 |

**Supplementary Table 8:** Random Forest pairwise strategy contrasts — p-factor. Columns as in Supplementary Table 2.

| Outcome | Contrast | Δz | SE (Δz) | t | df | 95% CI (z) | p-value |
|---|---|---|---|---|---|---|---|
| P-factor | Multi-Down − Local | -0.101 | 0.024 | -4.20 | 24 | [-0.151, -0.051] | 3.21e-04 |
| P-factor | Multi-Full − Local | -0.118 | 0.024 | -4.93 | 24 | [-0.168, -0.069] | 5.03e-05 |
| P-factor | Multi-Full − Multi-Down | -0.018 | 0.024 | -0.73 | 24 | [-0.067, 0.032] | 0.473 |

**Supplementary Table 9:** Site-level summary features and prediction accuracy (Random Forest). Per-site sample size, outcome dispersion SD(outcome), imaging-feature variance tr(Cov(X)), SD of the Multi-Full predicted values, and median Pearson r under each strategy across 100 splits.

| Outcome | Site | N | SD(outcome) | tr(Cov(X)) | SD(pred, MF) | Local r | Multi-Down r | Multi-Full r |
|---|---|---|---|---|---|---|---|---|
| Age | BHRC-1 | 128 | 1.988 | 4.35e+07 | 1.309 | 0.299 | 0.208 | 0.301 |
| Age | BHRC-2 | 292 | 1.758 | 4.46e+07 | 1.343 | 0.306 | 0.255 | 0.318 |
| Age | Colornest | 163 | 3.231 | 3.83e+07 | 1.250 | 0.177 | 0.057 | -0.006 |
| Age | HBNsiteCBIC | 540 | 3.555 | 4.76e+07 | 1.712 | 0.620 | 0.538 | 0.611 |
| Age | HBNsiteCUNY | 54 | 2.902 | 4.30e+07 | 1.416 | 0.086 | 0.114 | 0.432 |
| Age | HBNsiteRU | 438 | 3.494 | 4.62e+07 | 1.696 | 0.663 | 0.530 | 0.639 |
| Age | HBNsiteSI | 231 | 3.322 | 4.79e+07 | 1.704 | 0.506 | 0.448 | 0.535 |
| Age | NKI | 202 | 3.085 | 4.73e+07 | 1.707 | 0.618 | 0.434 | 0.590 |
| Age | PNC1 | 1,438 | 3.614 | 4.91e+07 | 1.862 | 0.606 | 0.530 | 0.554 |
| P-factor | BHRC-1 | 128 | 0.906 | 4.35e+07 | 0.112 | 0.016 | -0.044 | -0.149 |
| P-factor | BHRC-2 | 292 | 0.852 | 4.46e+07 | 0.096 | 0.042 | -0.024 | -0.126 |
| P-factor | Colornest | 163 | 0.827 | 3.83e+07 | 0.086 | -0.087 | -0.030 | -0.089 |
| P-factor | HBNsiteCBIC | 540 | 0.970 | 4.76e+07 | 0.109 | -0.077 | -0.014 | -0.026 |
| P-factor | HBNsiteCUNY | 54 | 0.875 | 4.30e+07 | 0.100 | 0.076 | -0.005 | -0.106 |
| P-factor | HBNsiteRU | 438 | 0.975 | 4.62e+07 | 0.146 | 0.063 | 0.021 | -0.016 |
| P-factor | HBNsiteSI | 231 | 1.018 | 4.79e+07 | 0.101 | 0.018 | -0.011 | 0.053 |
| P-factor | NKI | 202 | 0.875 | 4.73e+07 | 0.095 | 0.137 | 0.019 | 0.010 |
| P-factor | PNC1 | 1,438 | 0.940 | 4.91e+07 | 0.083 | 0.230 | 0.016 | 0.020 |

**Supplementary Table 10:** SHARP sensitivity analysis of training-strategy differences in prediction accuracy.

| Comparison | Δ*z* | 95% CI | Raw *p* | Holm *p* |
|---|---|---|---|---|
| **Ridge — Age** | | | | |
| Multi-Full − Local | 0.0561 | [0.0322, 0.0800] | 0.0003 | 0.0011 |
| Multi-Down − Local | −0.0914 | [−0.1096, −0.0732] | <0.0001 | <0.0001 |
| Multi-Full − Multi-Down | 0.1475 | [0.1338, 0.1613] | <0.0001 | <0.0001 |
| **Ridge — P-factor** | | | | |
| Multi-Full − Local | −0.0483 | [−0.0904, −0.0061] | 0.0291 | 0.0582 |
| Multi-Down − Local | −0.0663 | [−0.1027, −0.0299] | 0.0039 | 0.0117 |
| Multi-Full − Multi-Down | 0.0180 | [−0.0030, 0.0390] | 0.0700 | 0.0700 |
| **Random forest — Age** | | | | |
| Multi-Full − Local | −0.0022 | [−0.0256, 0.0212] | 0.8295 | 0.8295 |
| Multi-Down − Local | −0.1069 | [−0.1342, −0.0796] | <0.0001 | <0.0001 |
| Multi-Full − Multi-Down | 0.1047 | [0.0889, 0.1205] | <0.0001 | <0.0001 |
| **Random forest — P-factor** | | | | |
| Multi-Full − Local | −0.0933 | [−0.1358, −0.0508] | 0.0007 | 0.0028 |
| Multi-Down − Local | −0.0766 | [−0.1083, −0.0449] | 0.0011 | 0.0033 |
| Multi-Full − Multi-Down | −0.0167 | [−0.0465, 0.0131] | 0.1514 | 0.3028 |

*Note.* Δ*z* is the site-weighted mean Fisher-*z* difference across 120 half-experiments from 60 paired splits of nine sites. Positive differences favor the first strategy. Two-sided score tests and marginal 95% confidence intervals use the multisite SHARP working-model adaptation described in Supplementary Methods. Holm correction covers six comparisons per algorithm (three contrasts × two outcomes). Confidence intervals are not multiplicity-adjusted; significance is assessed using Holm $p < 0.05$.

**Supplementary Table 11:** Site-level summary features and prediction accuracy (Ridge). Per-site sample size, outcome dispersion SD(outcome), imaging-feature variance tr(Cov(X)), SD of the Multi-Full predicted values, and median Pearson r under each strategy across 100 splits.

| Outcome | Site | N | SD(outcome) | tr(Cov(X)) | SD(pred, MF) | Local r | Multi-Down r | Multi-Full r |
|---|---|---|---|---|---|---|---|---|
| Age | BHRC-1 | 128 | 1.988 | 4.35e+07 | 2.028 | 0.303 | 0.297 | 0.454 |
| Age | BHRC-2 | 292 | 1.758 | 4.46e+07 | 2.205 | 0.395 | 0.360 | 0.439 |
| Age | Colornest | 163 | 3.231 | 3.83e+07 | 1.969 | 0.222 | 0.052 | 0.113 |
| Age | HBNsiteCBIC | 540 | 3.555 | 4.76e+07 | 2.799 | 0.745 | 0.685 | 0.754 |
| Age | HBNsiteCUNY | 54 | 2.902 | 4.30e+07 | 2.149 | 0.447 | 0.313 | 0.715 |
| Age | HBNsiteRU | 438 | 3.494 | 4.62e+07 | 2.811 | 0.781 | 0.703 | 0.780 |
| Age | HBNsiteSI | 231 | 3.322 | 4.79e+07 | 2.460 | 0.688 | 0.591 | 0.717 |
| Age | NKI | 202 | 3.085 | 4.73e+07 | 2.515 | 0.668 | 0.576 | 0.728 |
| Age | PNC1 | 1,438 | 3.614 | 4.91e+07 | 2.616 | 0.679 | 0.647 | 0.670 |
| P-factor | BHRC-1 | 128 | 0.906 | 4.35e+07 | 0.095 | -0.031 | -0.002 | 0.016 |
| P-factor | BHRC-2 | 292 | 0.852 | 4.46e+07 | 0.098 | 0.065 | -0.021 | -0.012 |
| P-factor | Colornest | 163 | 0.827 | 3.83e+07 | 0.103 | -0.053 | 0.010 | -0.000 |
| P-factor | HBNsiteCBIC | 540 | 0.970 | 4.76e+07 | 0.103 | -0.121 | -0.006 | 0.026 |
| P-factor | HBNsiteCUNY | 54 | 0.875 | 4.30e+07 | 0.087 | -0.035 | -0.007 | 0.112 |
| P-factor | HBNsiteRU | 438 | 0.975 | 4.62e+07 | 0.119 | 0.108 | 0.039 | 0.103 |
| P-factor | HBNsiteSI | 231 | 1.018 | 4.79e+07 | 0.100 | 0.048 | 0.032 | 0.114 |
| P-factor | NKI | 202 | 0.875 | 4.73e+07 | 0.095 | 0.088 | -0.001 | 0.080 |
| P-factor | PNC1 | 1,438 | 0.940 | 4.91e+07 | 0.096 | 0.192 | 0.023 | 0.023 |

**Supplementary Table 12:** Ridge site-level regression — Multi-Full (n = 9 sites). Univariate regressions of each site's median Multi-Full Pearson r on four site-level features.

| Outcome | Strategy | Predictor | Slope | $R^2$ | p-value |
|---|---|---|---|---|---|
| Age | Multi-Full | SD(age) | 1.334e-01 | 0.168 | 0.273 |
| Age | Multi-Full | tr(Cov(X)) | 5.466e-08 | 0.683 | 0.006 |
| Age | Multi-Full | SD(predicted age, Multi-Full) | 5.377e-01 | 0.607 | 0.013 |
| Age | Multi-Full | Sample size | 1.319e-04 | 0.064 | 0.512 |
| P-factor | Multi-Full | SD(p-factor) | 3.895e-01 | 0.242 | 0.179 |
| P-factor | Multi-Full | tr(Cov(X)) | 4.859e-09 | 0.101 | 0.404 |
| P-factor | Multi-Full | SD(predicted p-factor, Multi-Full) | 4.082e-01 | 0.005 | 0.857 |
| P-factor | Multi-Full | Sample size | -2.796e-05 | 0.054 | 0.548 |

**Supplementary Table 13:** Ridge site-level regression — Local (n = 9 sites). As in Supplementary Table 12, for the Local strategy.

| Outcome | Strategy | Predictor | Slope | $R^2$ | p-value |
|---|---|---|---|---|---|
| Age | Local | SD(age) | 1.997e-01 | 0.425 | 0.057 |
| Age | Local | tr(Cov(X)) | 5.421e-08 | 0.758 | 0.002 |
| Age | Local | SD(predicted age, Local) | 2.531e-01 | 0.858 | 3.37e-04 |
| Age | Local | Sample size | 2.246e-04 | 0.209 | 0.216 |
| P-factor | Local | SD(p-factor) | 2.504e-01 | 0.028 | 0.669 |
| P-factor | Local | tr(Cov(X)) | 1.467e-08 | 0.255 | 0.166 |
| P-factor | Local | SD(predicted p-factor, Local) | 8.545e-01 | 0.135 | 0.330 |
| P-factor | Local | Sample size | 1.335e-04 | 0.339 | 0.100 |

**Supplementary Table 14:** Random Forest site-level regression — Multi-Full (n = 9 sites). As in Supplementary Table 12, for the Random-Forest Multi-Full model.

| Outcome | Strategy | Predictor | Slope | $R^2$ | p-value |
|---|---|---|---|---|---|
| Age | Multi-Full | SD(age) | 1.353e-01 | 0.194 | 0.235 |
| Age | Multi-Full | tr(Cov(X)) | 5.646e-08 | 0.818 | 8.03e-04 |
| Age | Multi-Full | SD(predicted age, Multi-Full) | 8.058e-01 | 0.754 | 0.002 |
| Age | Multi-Full | Sample size | 1.849e-04 | 0.141 | 0.319 |
| P-factor | Multi-Full | SD(p-factor) | 7.523e-01 | 0.453 | 0.047 |
| P-factor | Multi-Full | tr(Cov(X)) | 1.549e-08 | 0.518 | 0.029 |
| P-factor | Multi-Full | SD(predicted p-factor, Multi-Full) | 2.263e-03 | 0.000 | 0.999 |
| P-factor | Multi-Full | Sample size | 7.898e-05 | 0.216 | 0.207 |

**Supplementary Table 15:** Random Forest site-level regression — Local (n = 9 sites). As in Supplementary Table 12, for the Random-Forest Local model.

| Outcome | Strategy | Predictor | Slope | $R^2$ | p-value |
|---|---|---|---|---|---|
| Age | Local | SD(age) | 1.719e-01 | 0.287 | 0.137 |
| Age | Local | tr(Cov(X)) | 5.347e-08 | 0.673 | 0.007 |
| Age | Local | SD(predicted age, Local) | 3.430e-01 | 0.687 | 0.006 |
| Age | Local | Sample size | 2.775e-04 | 0.291 | 0.134 |
| P-factor | Local | SD(p-factor) | 1.065e-01 | 0.005 | 0.859 |
| P-factor | Local | tr(Cov(X)) | 1.637e-08 | 0.306 | 0.122 |
| P-factor | Local | SD(predicted p-factor, Local) | 1.128e+00 | 0.520 | 0.028 |
| P-factor | Local | Sample size | 1.337e-04 | 0.328 | 0.107 |

### Supplementary Figures

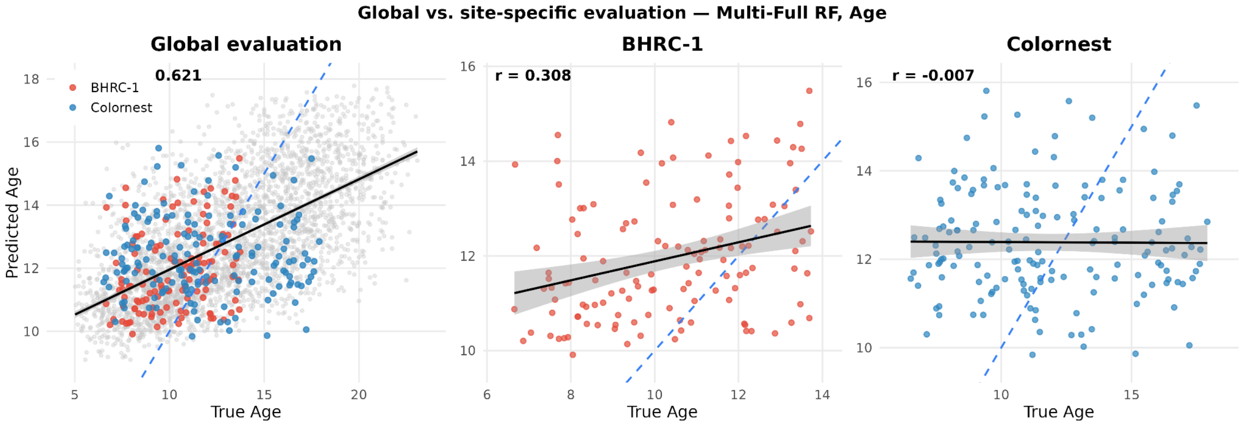


**Supplementary Figure 1:** Global vs. site-specific evaluation of the Multi-Full Random Forest model (Age). Left column: global scatter with one site highlighted; right column: that site alone.

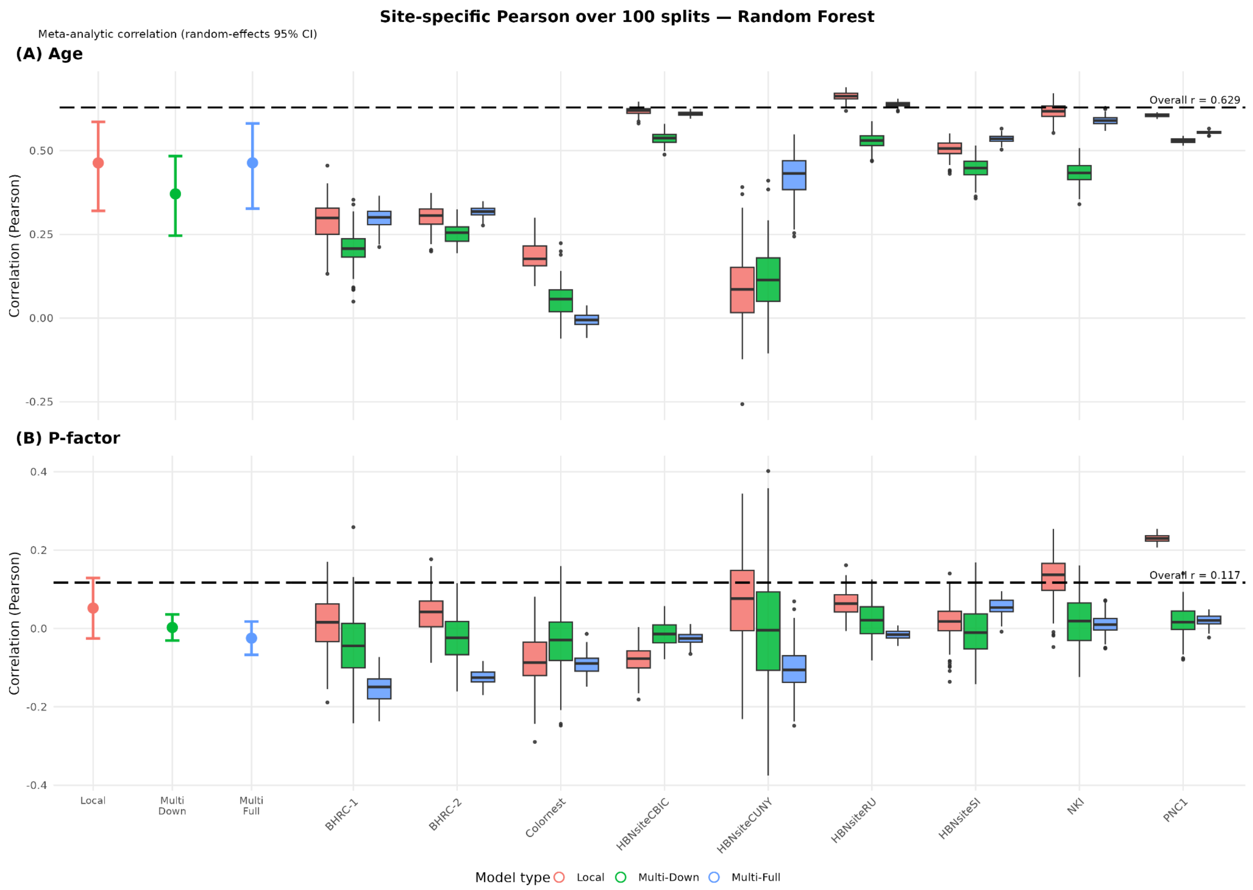


**Supplementary Figure 2:** Site-specific Pearson correlation over 100 splits (Random Forest). (A) Age, (B) P-factor. Left: pooled random-effects estimate with 95% CI per strategy. Right: per-site split distributions. Dashed line = Overall pooled r.

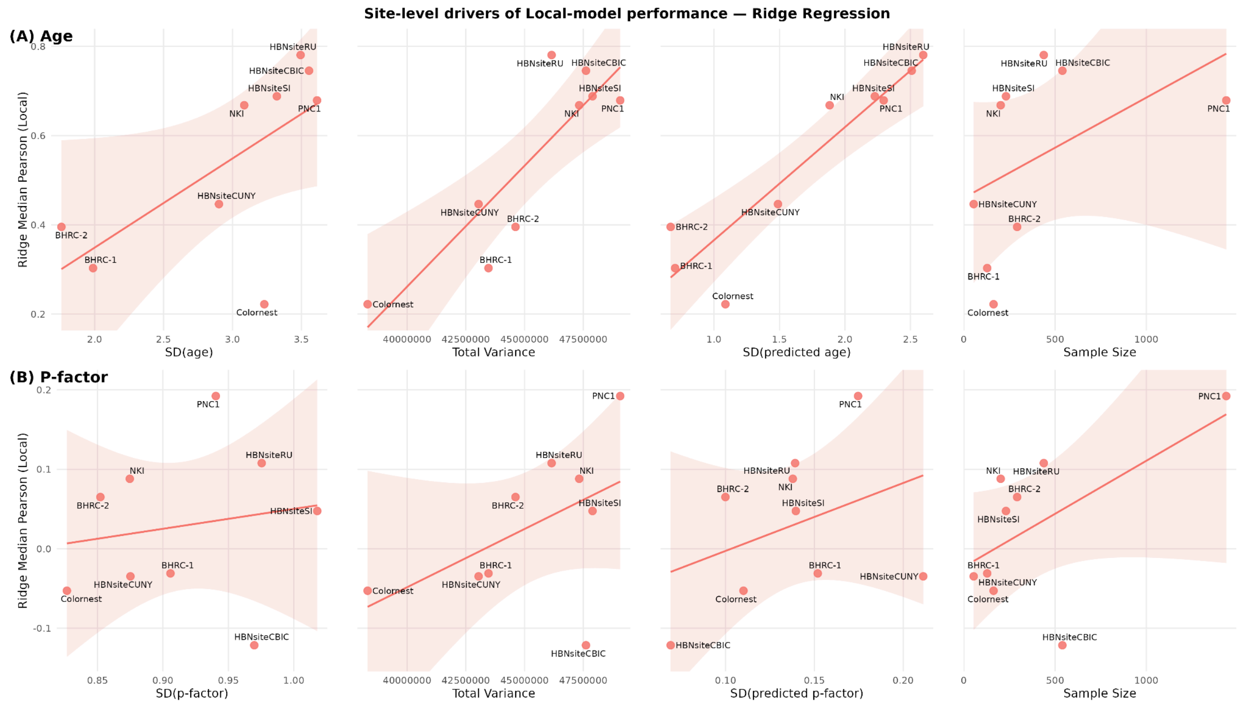


**Supplementary Figure 3:** Site-level drivers of Local-model performance (Ridge regression). Panels: SD(outcome), Total Variance, SD(predicted), and sample size, against median Local Pearson r. (A) Age, (B) P-factor.

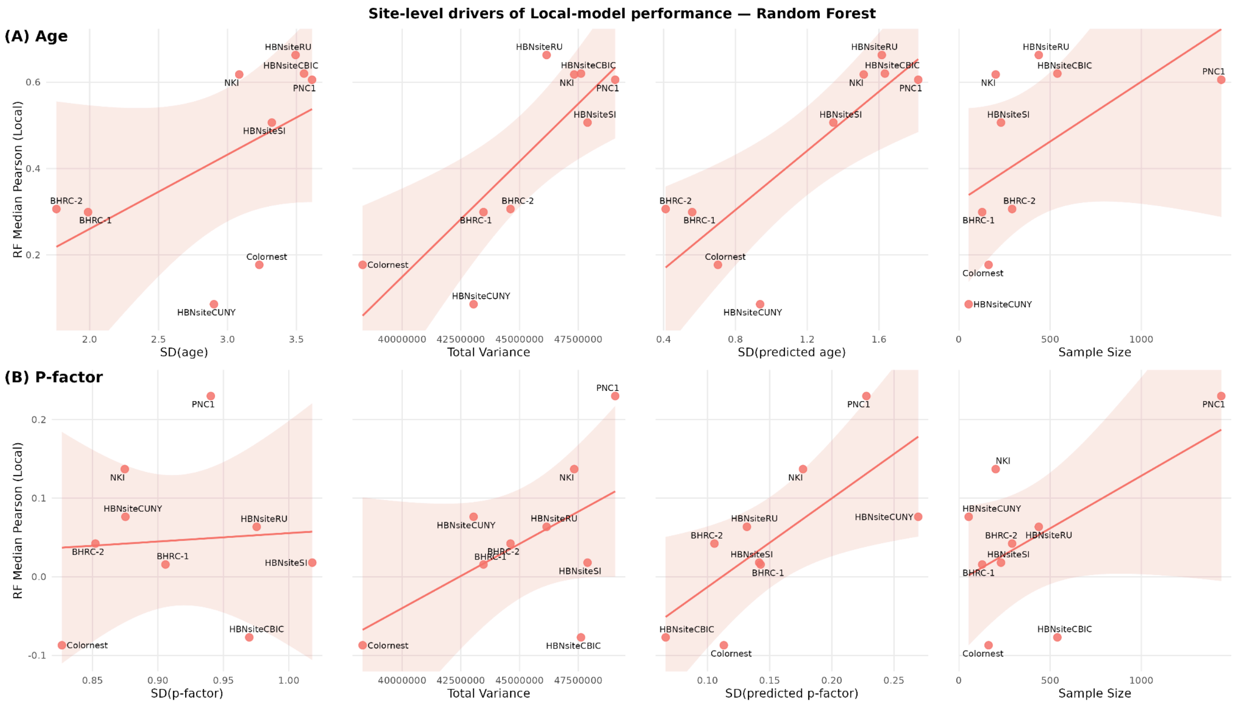


**Supplementary Figure 4:** Site-level drivers of Local-model performance (Random Forest). Panels: SD(outcome), Total Variance, SD(predicted), and sample size, against median Local Pearson r. (A) Age, (B) P-factor.

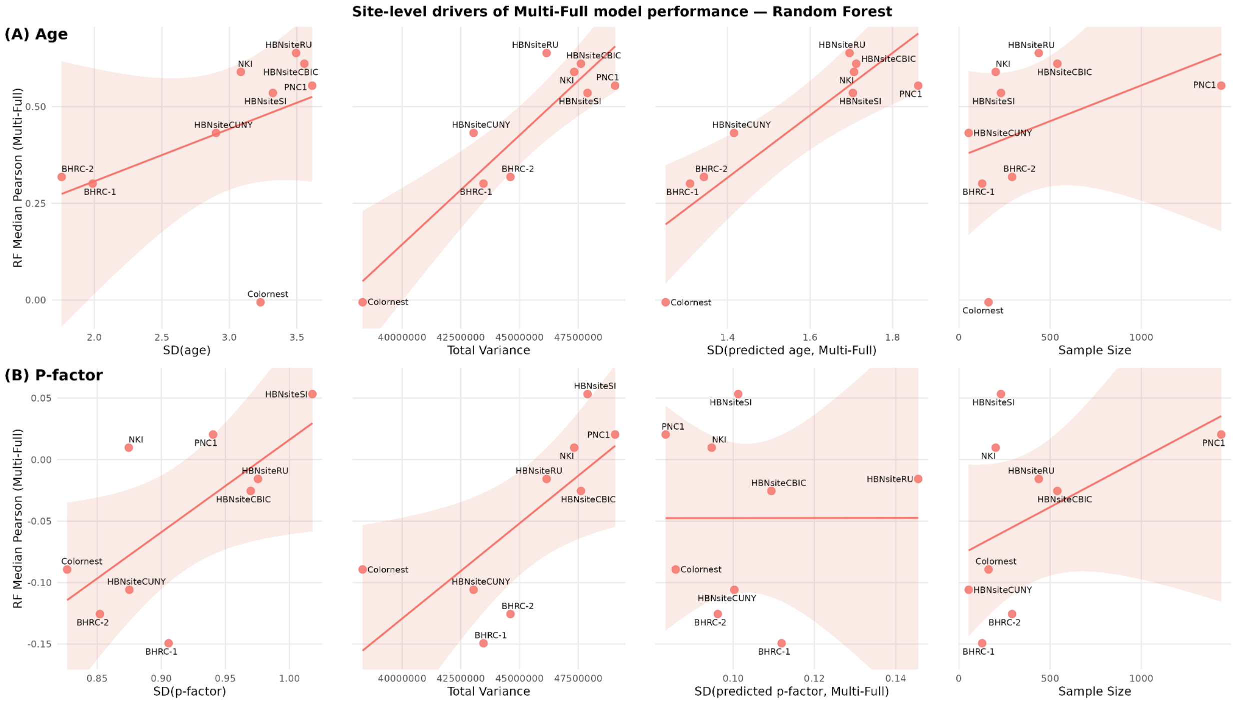


**Supplementary Figure 5:** Site-level drivers of Multi-Full-model performance (Random Forest). Same predictors as Supplementary Figure 3 against median Multi-Full Pearson r. (A) Age, (B) P-factor.
